# Vertical-junction Photodiodes for Smaller Pixels in Retinal Prostheses

**DOI:** 10.1101/2020.10.28.353128

**Authors:** Tiffany W Huang, Theodore I Kamins, Zhijie Charles Chen, Bing-Yi Wang, Mohajeet Bhuckory, Ludwig Galambos, Elton Ho, Tong Ling, Sean Afshar, Andrew Shin, Valentina Zuckerman, James S Harris, Keith Mathieson, Daniel Palanker

## Abstract

**Objective:** To restore central vision in patients with atrophic age-related macular degeneration, we replace the lost photoreceptors with photovoltaic pixels, which convert light into current and stimulate the secondary retinal neurons. Clinical trials demonstrated prosthetic acuity closely matching the sampling limit of the 100 μm pixels, and hence smaller pixels are required for improving visual acuity. However, with smaller flat bipolar pixels, the electric field penetration depth and the photodiode responsivity significantly decrease, making the device inefficient. Smaller pixels may be enabled (1) by increasing the diode responsivity using vertical p-n junctions and (2) by directing the electric field vertically using 3-D electrodes. Here, we demonstrate such novel photodiodes and test the retinal stimulation in a vertical electric field.

**Approach:** Arrays of silicon photodiodes of 55, 40, 30, and 20 μm in width, with vertical p-n junctions, were fabricated. The electric field in the retina was directed vertically by a common return electrode at the edge of the devices. Optical and electronic performance of the diodes was characterized in-vitro, and retinal stimulation threshold measured by recording the visually evoked potentials (VEPs) in rats with retinal degeneration.

**Main results:** The photodiodes exhibited sufficiently low dark current (<10 pA) and responsivity at 880 nm wavelength as high as 0.51 A/W, with 85% internal quantum efficiency, independent of pixel size. Field mapping in saline demonstrated uniformity of the pixel performance in the array. The full-field stimulation threshold was as low as 0.057±0.029 mW/mm^2^ with 10 ms pulses, independent of pixel size.

**Significance:** Photodiodes with vertical p-n junctions demonstrated excellent charge collection efficiency independent of pixel size, down to 20 μm. Vertically-oriented electric field provides a stimulation threshold that is independent of pixel size. These results are the first steps in validation of the feasibility of scaling down the photovoltaic pixels for subretinal stimulation.

## 1. Introduction

Atrophic Age-related Macular Degeneration (AMD) leads to loss of central vision due to the gradual demise of photoreceptors, a condition called geographic atrophy. This form of advanced AMD affects millions of patients: around 3% of people above the age of 75, and around 25% above 90 [1,2]. Currently, there is no therapy for such scotomata and the loss of sight is permanent. In these conditions, however, a significant number of the inner retinal neurons survive [3–6], providing an opportunity for functional restoration of vision by electrical stimulation of these neurons. Light-to-current conversion during phototransduction in photoreceptors includes a very strong amplification (up to six orders of magnitude: from a few photons per second to pA of current per cell). One approach to achieving this signal amplification for the electrical substitute of the photoreceptors is by electronic amplification in the subretinal implant [7]. However, since that requires a power cable, this approach is not suitable for AMD patients, where the preserved peripheral retina can be damaged by such a subretinal cable. For this purpose, we developed a wireless silicon photovoltaic subretinal implant activated by light [8]. Images of the visual scene captured by a video camera are processed and projected by augmented-reality glasses onto a subretinal photodiode array using intense pulsed light. Photovoltaic pixels in the array convert this light into biphasic pulses of electric current, which stimulate the retinal neurons in the inner nuclear layer (INL) – primarily the bipolar cells. To avoid perception of this light by the remaining photoreceptors, we use a near-infrared (∼880 nm) wavelength.

This system offers multiple advantages: (1) thousands of pixels in the implant can be activated simultaneously and independently; (2) since the pixels are activated by light, no wires are involved, which greatly simplifies the surgical procedure; (3) the external camera allows operation over a wide range of ambient illumination and provides user-adjustable image processing optimized for the dynamic range of the implant; (4) the optical nature of the implant maintains the natural link between eye movements and image perception; (5) network-mediated retinal stimulation retains many features of the natural signal processing, including antagonistic-center-surround [9], flicker fusion at high frequencies [10,11] and nonlinear summation of subunits, amongst others.

In prior studies, visual acuity (measured using alternating gratings in rodents) matched the 70 μm and 55 μm pixel size of the implants [11,12]. In the first clinical feasibility study of this technology, 5 patients with loss of central vision due to atrophic AMD have been implanted with a chip of 2 x 2 mm in size and 30 µm in thickness, containing 378 pixels, each 100 µm in size. All 5 patients perceive white-yellow prosthetic visual patterns with adjustable brightness in the previous scotomata, with prosthetic visual acuity in the best patient reaching 20/460 – only 10% below the level expected from the pixel pitch (20/420) [13]. Even more remarkable is that prosthetic central vision in AMD patients is perceived simultaneously with the remaining natural peripheral vision [14]. This success proves feasibility of the photovoltaic replacement of the lost photoreceptors for restoration of sight, and the main goal now is to improve its spatial resolution.

Since the loss of sight in AMD patients is limited by the central macula, visual acuity rarely drops below 20/400. Therefore, the majority of patients with atrophic AMD would benefit significantly if prosthetic acuity exceeds 20/100, which requires pixels smaller than 25 μm in size. However, simply scaling the pixel size down with flat arrays decreases the penetration depth of the electric field into the tissue, and therefore increases the stimulation threshold. With pixels smaller than 40 μm, the required charge injection exceeds the capacity of even one of the best electrode materials – sputtered iridium oxide (SIROF [15,16]).

When scaling photovoltaic pixels to smaller dimensions, their light-to-current conversion efficiency (photo-responsivity) also decreases with size, leading to further increase in the stimulation threshold. We activate the device by light at an 880 nm wavelength, and hence the Si thickness is about 30 μm – approximately matching the penetration depth of this radiation in Si. Traditionally, p-n junctions of photodiodes are made near the top surface (Figure 1a). A typical lifetime of the photo-generated carriers in lightly doped p-type silicon (10^15^ cm^-3^ acceptor density) is τ > 10 μs [17]. Before recombining, photogenerated carriers can diffuse over a distance L ∼ (D× τ)^0.5^ > 200 μm, which greatly exceeds the pixel pitch (Diffusion coefficient D ∼ 35 cm^2^/s for electrons and ∼12 cm^2^/s for holes [18,19]) and would lead to sharing of charge between pixels. To prevent this spread of charges into the neighboring pixels, the diodes should be separated by boundaries that prevent photogenerated carriers from moving from one pixel to adjacent pixels. In our previous arrays, pixels were separated by full-depth trenches with oxidized sidewalls and filled with polysilicon, as shown in Figure 1a [12,20].

**Figure 1.**
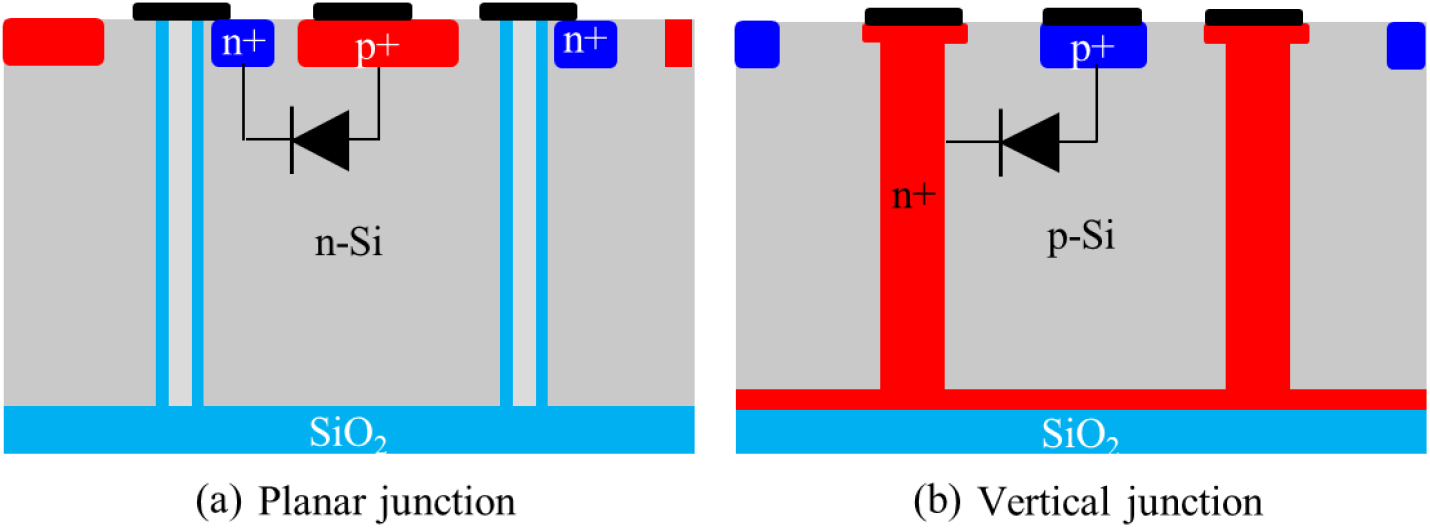
Diagram of the photovoltaic pixel with (a) planar and (b) vertical junctions. The dark blue region represents the highly doped area of the same polarity as the background doping for making ohmic contact, while red has the opposite doping polarity. The p^+^ region is connected to the active electrode, while the n^+^ - to the return electrode.

However, the vertical reactive ion etch of the silicon creates significant crystal damage in the adjacent areas, resulting in traps and mid-gap recombination centers for carriers [21–24], and the finite surface recombination velocity at the Si/SiO_2_ interfaces at the diode sidewalls also causes loss of photogenerated carriers, As the size of the photosensitive region in the pixel becomes smaller, a greater fraction of the photogenerated carriers recombine at the sidewalls of the pixel, precluding their collection at the p-n junction. Such recombination results in decreasing responsivity with reduced pixel size: 0.24, 0.22, and 0.20 A/W with pixel sizes of 70, 55, and 40 µm, respectively [16]. It is important to keep in mind that in the previous design, pixels were composed of two diodes connected in series, and therefore the diode width was less than half of the pixel size. Thermal oxidation of the sidewalls helps mitigate the effect of some of the etch-induced defects, as it results in some consumption of the defective silicon caused by etching [25]. However, thicker oxide on the side walls decreases the photosensitive area of the photodiodes. In addition, with smaller pixels, the density of the isolation trenches increases, leading to further loss of photosensitive area of the device and resulting in oxidation-related stress in the wafer [26], which may warp the wafer beyond the curvature suitable for lithographic processing.

To address this problem, we now placed vertical p-n junctions near the isolation trenches, thereby eliminating the oxidation-related stress in Si, minimizing recombination loss and maximizing carrier collection (Figure 1b). Vertical p-n junctions have been proposed since the 1970s [27] and allow decoupling photon absorption from carrier collection. This approach allows designers to choose a thickness of silicon to properly absorb the wavelength of interest, while independently setting the required electrode spacing for efficient charge collection. Such structures have been used for high-intensity solar cells [28] and high-energy photodetectors [29–31]. To eliminate the effect of interface recombination at the Si-SiO_2_ interface at the bottom of the pixel, we implanted phosphorus into the p-type device wafer before the device wafer and handle wafer were bonded to form the silicon-on-oxide (SOI) structure. This buried n^+^ layer creates a p-n junction near the back surface of the device layer so that minority carriers diffusing to that surface are also collected.

To resolve the problem with a limited penetration of electric field into tissue in front of small pixels, we proposed a novel 3-dimensional structure above the photodiode array to interface with the retina, which we call a “honeycomb” [16]. This approach is based on migration of the retinal cells into cavities in the subretinal implant [32,33] (Supplementary Figure 1). Wells of the honeycomb, with an active electrode at the bottom and a return electrode on top, direct the electric field vertically to match the vertical orientation of bipolar cells (Figure 2b,c). This enhances the cellular polarization, and thereby reduces the stimulation threshold. Walls of the honeycomb provide electrical isolation of the stimulation zone within each pixel, thereby enabling nearly 100% spatial contrast of the stimulation patterns, i.e. practically no crosstalk with the neighboring pixels. Moreover, this geometry decouples penetration depth of the electric field from the pixel width, thus enabling a nearly constant stimulation threshold in terms of irradiance and current density for smaller pixels [16].

**Figure 2.**
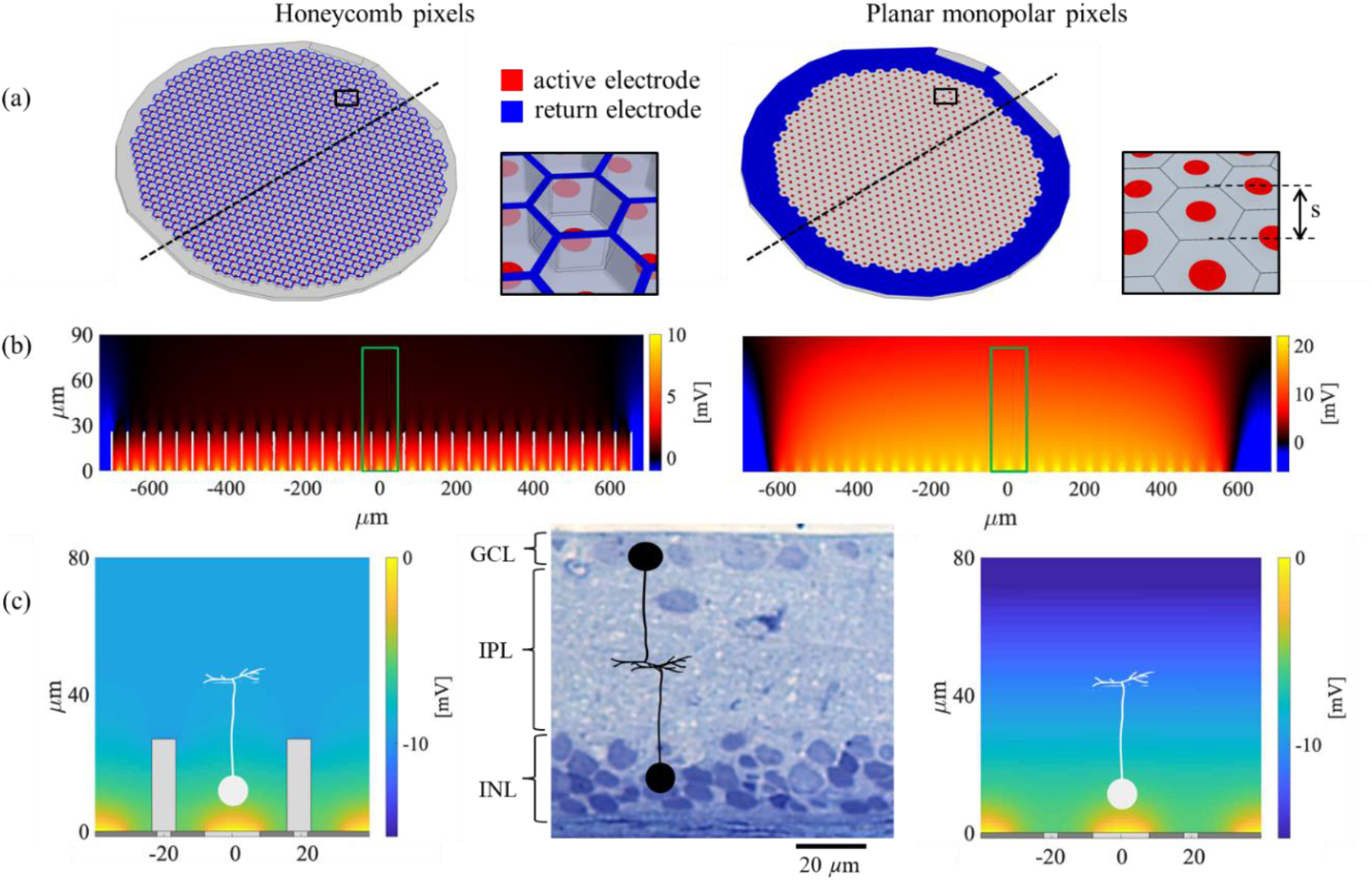
(a) Layout of the photovoltaic arrays with 40 μm pixels, having a common return electrode on top of the honeycomb walls (denoted ‘Honeycomb pixels’ in the left column) and in the periphery of the device (denoted ‘Planar monopolar pixels’ in the right column). The size of the pixel is denoted with ‘s’. (b) Electric potential in the medium relative to infinity produced by full-field illumination, with electric current of 100 nA per pixel. Note different scale in vertical and horizontal directions, introduced for better visibility. (c) Zoom into the single pixel area outlined in (b) (green box). In this case, the horizontal and vertical scales are the same, and potential is plotted relative to the center of the active electrode right above the device, showing similar voltage drop across the bipolar cell for these two electrode geometries. Histology of the degenerate retina (RCS) with a diagram of the bipolar and ganglion cells (BC and RGC) is shown to scale for comparison. Confocal microscopy of the retinal cells migrating into the honeycombs is shown in Supplemental Figure 1.

Since fabrication of the honeycomb structures with a return electrode on top requires significant development, in this paper we experimentally evaluate performance of the photovoltaic array with vertical orientation of electric field but using planar electrodes, as a first step toward 3-D arrays. To create nearly vertical electric field within the retinal thickness and thereby make the retinal stimulation threshold of such a planar array similar to that expected with honeycombs, we placed the common return electrode on the periphery of the device (Figure 2a). Simultaneous activation of all the pixels in such array (full-field illumination) creates a nearly vertical electric field within the thickness of the retina, which is similar to the field within the honeycombs. Due to deeper penetration of the vertical electric field without the local returns, the voltage drop across the bipolar cells in a uniform field is about two times larger than with honeycombs (Figure 2c), and hence the full-field retinal stimulation threshold is expected to be a bit lower [34]: Based on the previously described modeling [16], for the honeycomb pixels having an active electrode diameter of 40% the pixel size, stimulation threshold with 10 ms pulses is about 0.2 mA/mm^2^ on the active electrodes. With electrodes of the same size, but the return electrode at the edge of the implant, stimulation threshold is expected to be about 0.12 mA/mm^2^. Due to high crosstalk between the neighboring pixels in front of such a monopolar array, spatial resolution is not expected to be close to the size of a single pixel. Therefore, in the current study, we focus on validation of the stimulation threshold in a vertical field (full-field stimulation), while spatial resolution will be assessed in the future using arrays with integrated honeycomb return electrodes.

To summarize, in this article we describe the modeling, fabrication, and evaluation of silicon pixels with vertical p-n junctions for a subretinal prosthesis. We examine (1) the enhancement in the light responsivity of smaller pixels with vertical p-n junctions and (2) the improvements in stimulation threshold with a vertical electric field.

## 2. Materials and Methods

### 2.1 Photovoltaic array design and fabrication flow

Silicon is the material of choice for the photovoltaic retinal implant due to its high quantum efficiency, sensitivity to near-infrared light, and reliability of the fabrication procedures. The light intensity required for photovoltaic stimulation is on the order of 1 mW/mm^2^ [12,16], which is about 1000 times brighter than the upper end of natural illumination on the retina. Therefore, for restoration of sight in patients who retain some light sensitivity, the light activating the photovoltaic pixels should be in the invisible part of the spectrum. As shown in Figure 3a, photoreceptor sensitivity drops from its peak by about 6 orders of magnitude at 800 nm, and by about 7 orders of magnitude around 880 nm. On the other hand, ocular transmittance starts rapidly decreasing beyond 900 nm due to water absorption, while it is still above 80% near 880 nm. Typical silicon photodiode responsivity also peaks near 880 nm [35], while the radiation penetration depth at this wavelength is around 25 μm, as shown in Figure 3b. Therefore, we design our system for 880 nm wavelength and 30 μm device layer thickness, which provides about 90% absorption when the back surface is metalized (double pass).

**Figure 3.**
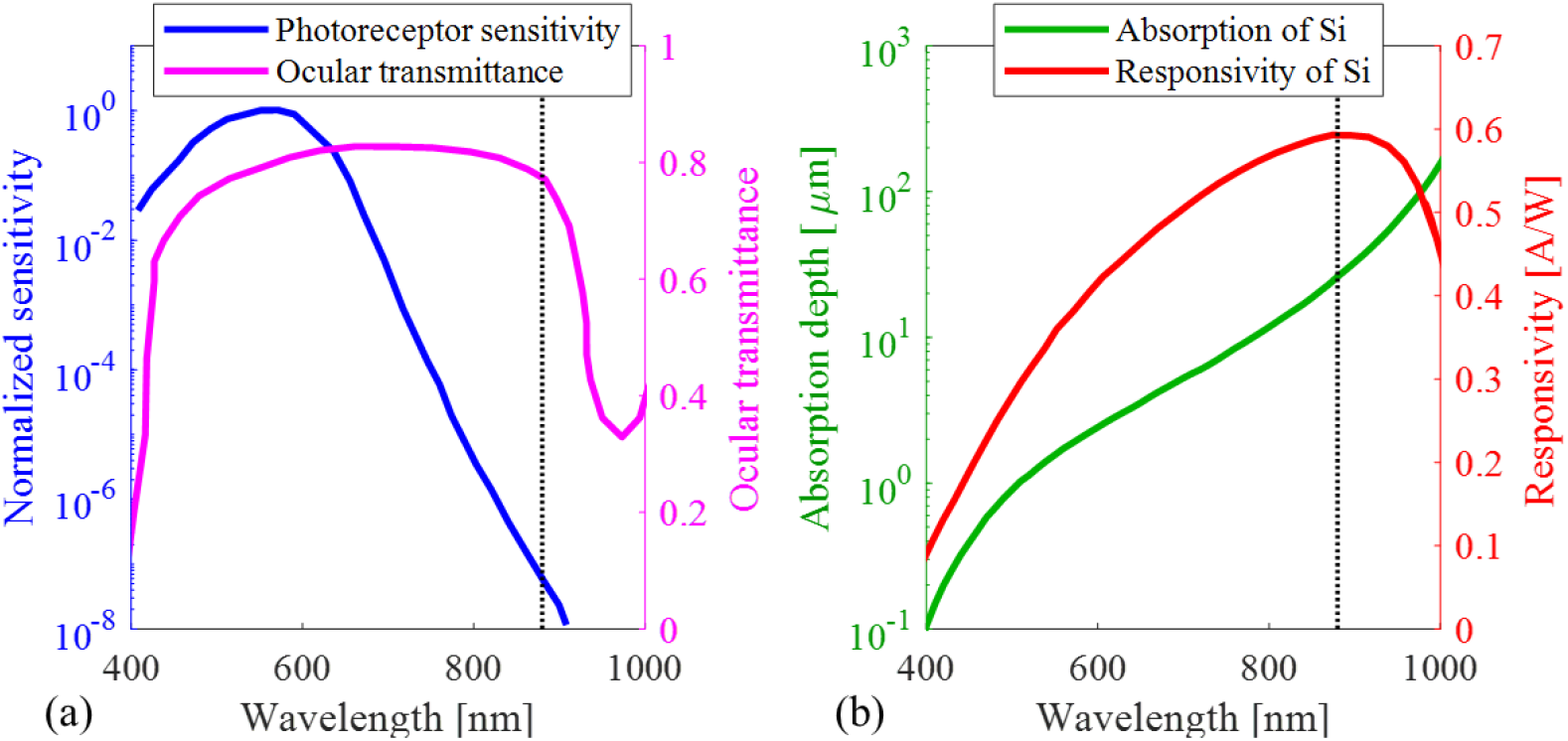
(a) Normalized photoreceptor sensitivity[36] and ocular transmittance to the retina [37] as a function of wavelength, showing high transmittance and low photoreceptor sensitivity near 880 nm wavelength. (b) Silicon absorption length, and typical silicon photodiode responsivity(A/W)[35] as a function of wavelength, showing absorption depth around 25 µm and high responsivity near 880 nm wavelength.

Our photodiode arrays with hexagonal pixels are designed with sizes of 55, 40, 30, and 20 µm, as defined in Figure 1a, on lightly boron-doped silicon-on-insulator SOI wafers with a buried n^+^ layer ion implanted (phosphorus with a dose of 2×10^15^ cm^-2^ at 80 keV) adjacent to the buried oxide before bonding to form the SOI wafer. Figure depicts the array fabrication process, which consists of eight mask layers: forming the vertical junction (a-d), forming the ohmic contact to the electrodes (e-f), forming the anti-reflection coatings and metal electrodes (g-i), adding the high-capacitance coating on the electrodes and releasing the device (j-l). Following the devices’ release, they are flipped over, and the sidewalls and back of the devices are coated with titanium (not shown in a figure).

### 2.2 Vertical p-n junctions

Trenches used to form the vertical p-n junctions are designed with a width of 1.7 μm, narrow enough for the junction doping to fit under the metal electrodes and wide enough for filling and for dopant diffusion. A thermal oxide of 1.5 μm on the top surface of the wafer provides a hard mask for fabrication of the trenches used for the vertical junction; the oxide is then used as a mask for etching the 30 μm thick silicon layer. For ease of filling, we design a slight tilt of 0.5° in the trench walls. The top silicon oxide thickness is designed to ensure sufficient protection of the silicon despite the oxide loss during the subsequent fabrication steps. The deep reactive ion etch (DRIE) of Si is performed at the University of Michigan fabrication facility, followed by gas-phase dopant diffusion using phosphorus oxychloride (POCl_3_) at the University of California, Berkeley. Subsequently, a short calibrated buffered-oxide etch (BOE) removes the brittle phosphosilicate glass (PSG) formed during POCl_3_diffusion, but not the buried oxide. LPCVD-deposited polysilicon fills the trenches, which are then planarized by chemical mechanical polishing. To prevent excessive wafer curvature caused by the processing-induced stress, similar trenches are created and filled on the back side of the wafer.

SEM images (Figure 5a) confirm the uniformity of the etch, with less than 100 nm variation in the trench width from center to edge of the wafer, as well as the consistency from wafer to wafer and minimal notching adjacent to the buried oxide. Figure 5b shows the cross-section of the device after the side walls are doped and trenches filled, demonstrating the resulting depletion zone (*) along the sides and bottom of the device, visible due to effectively charging the device during electron microscopy. Simulation using TCAD Sentaurus suggests that the dopant diffusion and subsequent heat treatment parameters should result in sufficient n^+^ doping within the polysilicon trench filling for high carrier density and high conductance in the trenches and adjacent single-crystal silicon. Nanoscale secondary ion mass spectroscopy (SIMS) confirms that the dopant diffusion is consistent and uniform across the trench (Figure 5c). The SIMS preparation involves embedding the sample in epoxy and polishing for flatness, which results in some irregularities at the top surface, as seen in the inset SEM. The SIMS curve is normalized by dividing the phosphorus by the silicon SIMS values, integrating along the trench, and then shifting the peak to the solid solubility of phosphorus in silicon at the highest annealing temperature in the fabrication process.

**Figure 4.**
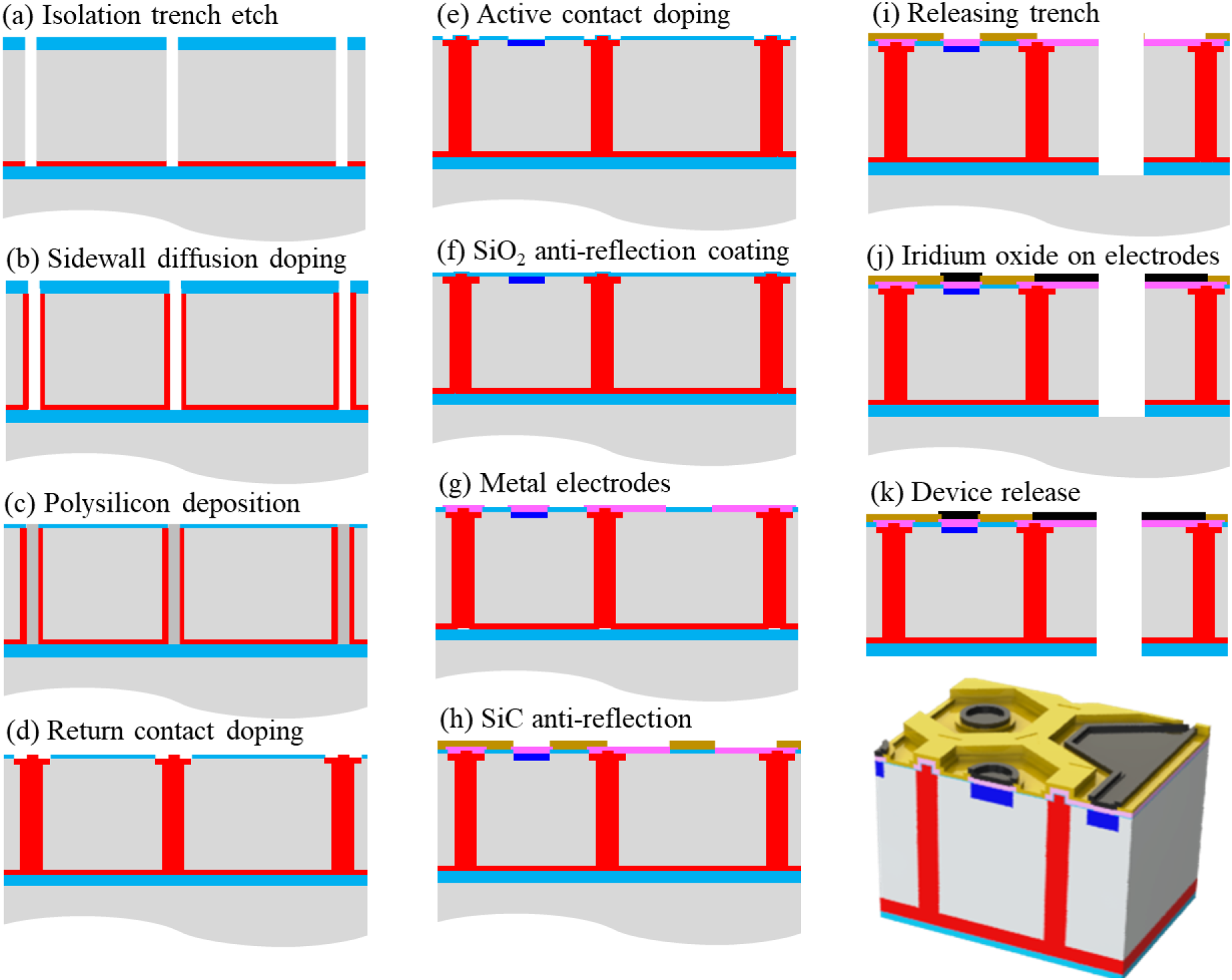
Fabrication process flow for photodiodes with vertical p-n junctions. Layers shown not to scale for better visibility.

**Figure 5.**
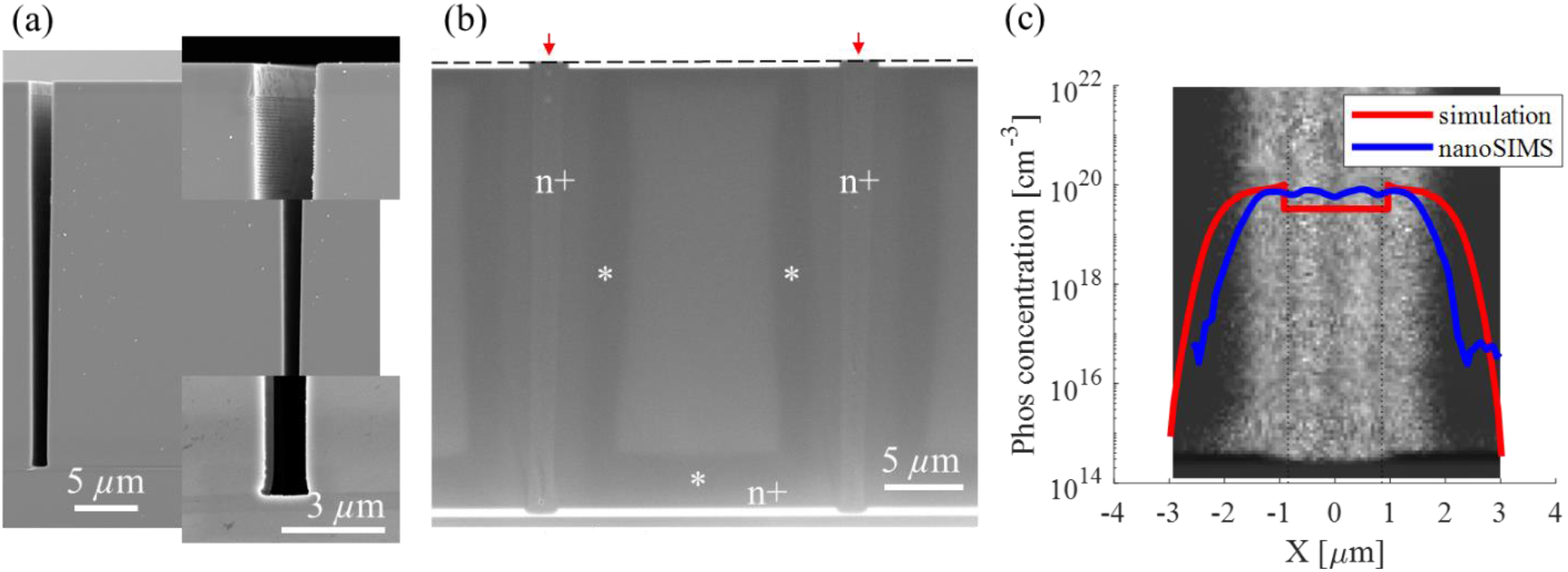
(a) Trenches of 1.7μm on top and 1.2μm at the bottom, etched for pixel isolation and p-n junction formation. Magnified top and bottom sections are shown on the right. (b) Cross-section of the device with trenches filled by polycrystalline silicon (pointed by the red arrows). The area adjacent to the trench is n-doped, and the adjacent dark stripes are the depletion zones (*). Dash line shows the top of the SiO_2_ layer not visible due to the white background. (c) SIMS profile of dopant concentration across the vertical junction overlying the scanning electron microscope (SEM) image, confirming consistent and uniform doping across the trench.

### 2.3 Ohmic contact formation

The n^+^ contacts to the junction are formed by first etching through the oxide hard mask, then doing a subsequent POCl_3_ diffusion to achieve ∼4 Ω /sq sheet resistance. The contacts to the p-type bulk silicon are formed by a subsequent oxidation, etching through the oxide hard mask, growing a screening oxide to prevent damage near the surface and ion implanting boron (Innovion Corporation, San Jose, CA) at 20 keV with a dose of 4 x 10^15^ cm^-2^. The doping must be high enough to ensure that the contacts are ohmic, not Schottky, and that they do not cause major power dissipation or voltage drop. The achieved contact resistivity, measured using a Kelvin structure, is quite reasonable: 1.2 x 10^−7^ Ω ·cm^2^ and 2.1 x 10^−6^ Ω ·cm^2^ for n-well and p-well, respectively. The expected power dissipation, calculated by multiplying the square of the current by the contact resistance, is below 0.04% of the power generated by the photodiode, assuming the maximum (open circuit) voltage generation.

### 2.4 Anti-reflection coating and electrode formation

To minimize the loss of light due to reflection from the device surface, we use an anti-reflection coating consisting of SiC and SiO_2_ (Figure 6a). To preclude pinholes, and to ensure electrical insulation, the SiC layer is designed to be greater than 150 nm thick, and the underlying SiO_2_ layer should be at least 55 nm thick. We use the transfer-matrix method to optimize the anti-reflection layer thicknesses for incident light at 880 nm, averaging from 878 to 882 nm to account for the 4 nm bandwidth of our laser. We find that at 55 nm SiO_2_ and 208 nm SiC, the reflectance of light from the top surface is about 2.4% (Figure 6b). About 73% of the light entering silicon will be absorbed in the 30 µm thick device layer of silicon. Of the light transmitted through silicon, about 70% will be reflected back from the metalized bottom of the device, and another 73% of that will be absorbed in Si on the second pass, adding up to about 85% of the incident light absorbed in Si, 7% reflected from the device, and 8% absorbed in Ti. Since the coherence length (*L* = λ^2^/(2π*n*Δλ)) of the λ = 880 nm light with Δλ = 4 nm bandwidth in Si (*n* = 3.6) is about *L* = 9 μm, the light reflected from the back of the device is incoherent with that entering from the front, and therefore their intensities are added as scalars without taking into account the phase.

**Figure 6.**
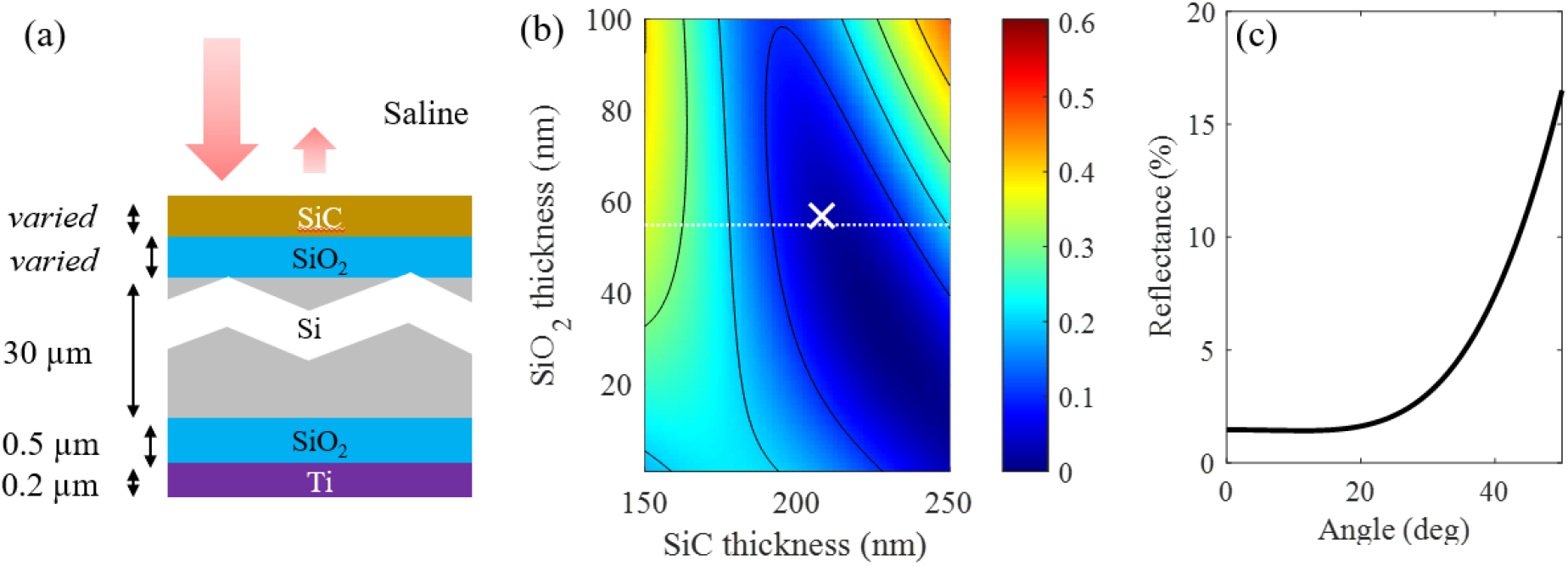
(a) Diagram of the stack used to design the anti-reflection coating. The thicknesses of the silicon, buried silicon oxide, and titanium are 30, 0.5, and 0.2 µm, respectively. The silicon carbide and top silicon oxide thicknesses are varied. (b) Reflectance of the front surface, as a function of SiC and SiO_2_ thickness, calculated by the transfer matrix method. White dash line indicates a safe SiO_2_ thickness for avoiding pinholes. The ‘X’ marks our attained layers of 57 nm SiO_2_ and 208 nm SiC, achieving 2.4% reflectance. (c) Reflectivity as a function of the incidence angle.

As can be seen in Figure 6c, reflectivity remains very low until the incidence angle reaches about 20°. Light incidence angle on the central macula in a human eye does not exceed 10° (pupil radius <3mm, distance to the retina >20mm), while in a dilated rat eye it can reach up to 20°. Therefore, reflectivity should stay sufficiently low for the incidence angles accessible in the eye.

To form the anti-reflection coatings and the electrodes, we first strip all oxide in a 6:1 buffered oxide etch, perform dry oxidation at 1000°C to form thermal oxide for the first coating, and then use bilayer lithography to define the Ti/Pt metal stack to form the active and return electrodes using a lift-off process. Then we deposit amorphous SiC using PECVD (EIC Biomedical, Norwood, MA) at 325°C. This SiC also functions as a protective coating [38]. We achieved 57 nm SiO_2_ and 208 nm SiC (shown in Figure 6b by the ×), which provides 2.4% reflectance. Next, we sequentially etch the SiC vias and the releasing trenches, and finally, we use lithography and a lift-off process to define the sputtered iridium oxide (SIROF) (deposited at EIC Biomedical, Norwood, MA) layer on the electrodes.

### 2.5 Release of the photodiode arrays

For the release from the wafer, we attached the devices to a carrier wafer on the front surface using a thick protective spray-coated resist, and then used grinding to thin the handle wafer from the back to 50 μm, followed by xenon difluoride etching to remove the remainder of the handle wafer and expose the buried oxide layer. For the final release of the arrays from the carrier wafer, the photoresist is dissolved in acetone and isopropanol. The released devices are then mounted face-side down on a wafer coated with photoresist and sputtered with 200 nm titanium to coat the sides and back of the devices. To remove the residue of the photoresist from the top surface of the device and, most importantly, from the porous SIROF coating of the electrodes, the devices then are cleaned with N-methyl-pyrrolidine (NMP) based solution (Shipley Microposit Remover 1165) and finally, with sodium hypochlorite (NaClO). The released devices are shown in Figure 7.

**Figure 7.**
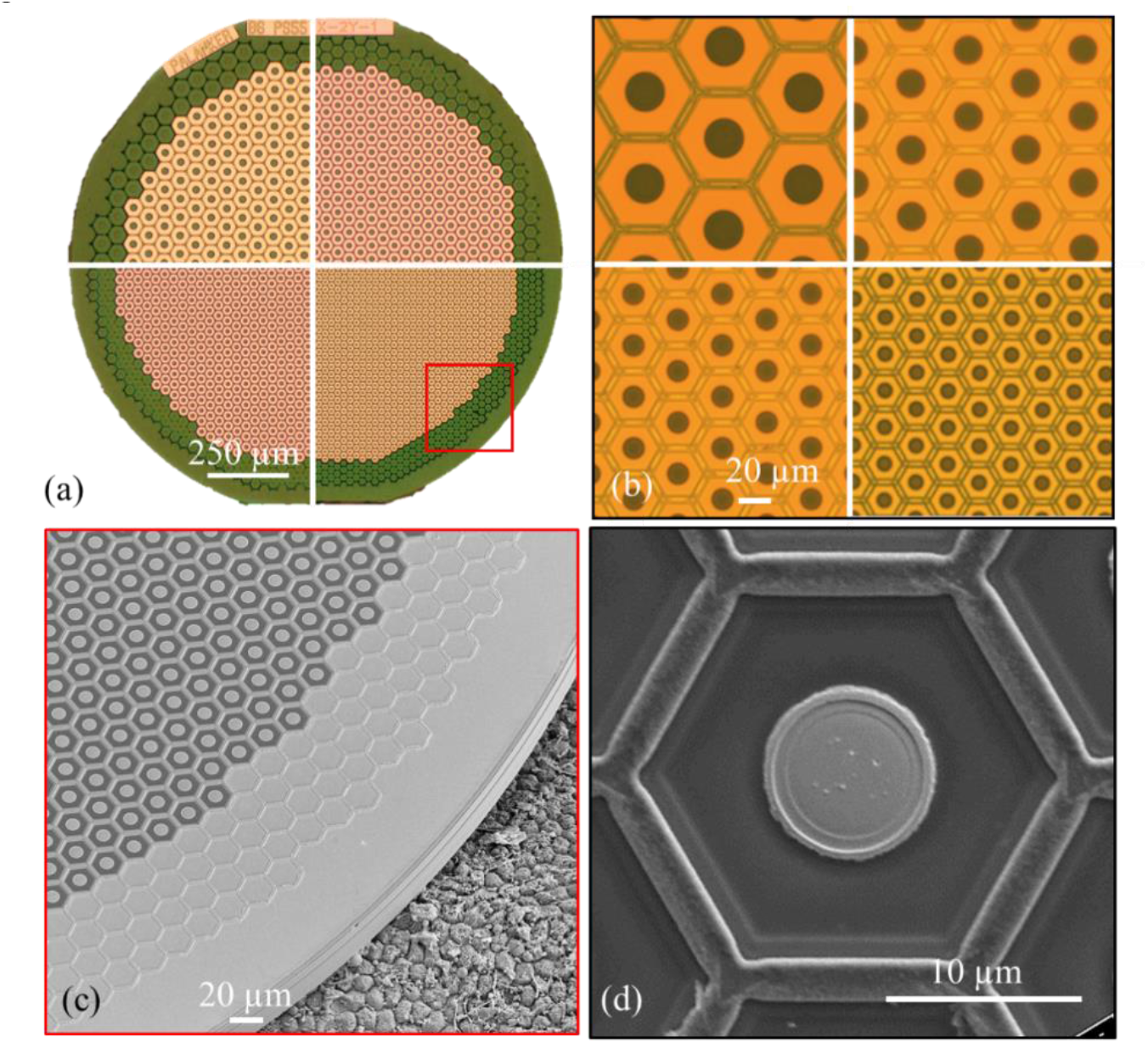
(a-b) Optical images showing 1.5 mm implants with pixel size 55, 40, 30, and 20 µm (cropped from 4 different arrays). (c) SEM image of the released device with 20 µm pixels, placed on a porcine retinal pigment epithelium for scale, and (d) SEM of a single 20 µm pixel, showing a central active electrode, surrounded by photosensitive area, and the hexagonal trench covered by metal buried under SiC to connect to the outer return electrode ring.

### 2.6 Ex-vivo characterization of the device

The I-V curve of the fabricated diodes is measured by sweeping the voltage from −1 to 0.8 V across the diode and measuring the corresponding current, limited by 0.6 mA compliance to avoid thermal damage. The light-to-current conversion is measured with an 880 nm laser (JOLD-30-FC-12, Jenoptik, Jena, Germany) illuminating the diode at irradiances varying between 2 and 8 mW/mm^2^, and the photosensitive area for this conversion is calculated from focused-ion beam (FIB)SEM images.

The diffusion capacitance, associated with the charge storage in the quasi-neutral region of the p-n junction, is measured using electrical impedance spectroscopy. These measurements are performed with a potentiostat (Interface1010E, Gamry Instruments, Warminster, PA) using the two-electrode setup, with 10 mV-RMS perturbation on top of the bias voltage, and at 10 frequencies per decade in the range between 10 Hz and 1 MHz. The working and the counter electrodes are connected to the p and the n ends of the photodiode, and measurements are performed in the dark. The bias voltage is set at 12 different levels – every 0.1 V between 0 and 0.5 V, and every 0.05 V between 0.5 and 0.8 V.

The quality of the SIROF electrodes is characterized using impedance spectroscopy on test structures, with the same equipment and at similar voltage settings. The capacitance values are then extracted by fitting the spectra with the corresponding circuit models. The electrode-electrolyte interfaces of SIROF are often modeled by a Randles model [39], the capacitor of which is indicative of the charge injection capacity (CIC)[40,41]. More details of the fitting and the electrochemical cell are described in the Supplementary Material.

To check uniformity of the photovoltaic conversion in pixels and of SIROF coating on electrodes, the electrical signals from each pixel are mapped in electrolyte under uniform full-field illumination at 880 nm, pulsed at 3 Hz. Devices with 20 and 40 μm pixels were placed in a diluted phosphate buffered saline (PBS) solution, with resistivity of 570 Ω ·cm, matching the expected retinal resistivity [34]. The potential at the plane 20 μm above the devices in the medium is recorded by scanning a micro-pipette in steps of 20 μm, when the devices are illuminated by 5 ms laser pulses of 5 mW/mm^2^ at 3Hz repetition rate [42].

### 2.7 Electrode design and modeling

The electric field generated by the monopolar device in electrolyte is calculated with a 3-D finite element model (FEM) of a complete array of 40 *μ*m pixels using COMSOL Multiphysics 5.4. The modeled device, 1.5 mm in diameter and 30 *μ*m in thickness, consists of over one thousand 40 *μ*m hexagonal pixels, each of which has an active electrode 16 *μ*m in diameter. The electrostatic module in COMSOL solves Poisson’s equation for electrical conduction, assuming a steady-state electric current, with the model parameters described in our previous publications [34]. The threshold value of the network-mediated retinal stimulation, defined by the visually evoked potential (VEP), is estimated as described in our previous publication [16].

### 2.8 Surgical procedures and VEP measurements

All experimental procedures are conducted in accordance with the Statement for the Use of Animals in Ophthalmic and Vision research of the Association for Research in Vision and Ophthalmology (ARVO), and approved by the Stanford Administrative Panel on Laboratory Animal Care. Royal College of Surgeons (RCS) rats are used as an animal model of the inherited retinal degeneration. The RCS colony is maintained at the Stanford Animal Facility under 12h light/12h dark cycles with food and water *ad libitum*. The devices are implanted subretinally, as previously described [10,11], after a complete loss of the outer nuclear layer evidenced by optical coherence tomography (OCT; HRA2-Spectralis; Heidelberg Engineering, Heidelberg, Germany). A total of 8 animals were implanted with 1.5 mm diameter arrays containing pixels of 40 µm (n=4) and 20 µm (n=4). Animals are anesthetized with a mixture of ketamine (75mg/kg) and xylazine (5mg/kg) injected intraperitoneally. A 2 mm incision is made through the sclera and choroid 1.5 mm posterior to the limbus. The retina and RPE are separated with an injection of saline solution into the subretinal space, and the implant is inserted. The conjunctiva is sutured with nylon 10-0, and topical antibiotic (bacitracin/polymyxinn B) is applied on the eye postoperatively. The animals are monitored using OCT to visualize the retina and the implant over time.

For measurement of the visually evoked potentials (VEP), each animal was implanted with three transcranial screw electrodes: 1 electrode over each hemisphere of the visual cortex (4 mm lateral from midline, 6 mm caudal to bregma), and a reference electrode (2 mm right of midline and 2 mm anterior to bregma). Following anesthesia and pupil dilation, the cornea of the animal is covered with a viscoelastic gel and a cover slip to cancel the cornea optical power and ensuring good retinal visibility. The retinal implant is illuminated with a customized projection system, consisting of a near-infrared laser at 880 nm (MF_880nm_400um, DILAS, Tucson, AZ), collimating optics, and a digital micromirror display (DMD; DLP Light Commander; LOGIC PD, Carlsbad, CA) for patterning. The entire optical system is integrated with a slit lamp (Zeiss SL-120; Carl Zeiss, Thornwood, NY) for convenience of observing the illuminated retina via a CCD camera (acA1300-60gmNIR; Basler, Ahrensburg, Germany).

For the stimulation threshold measurements, the NIR illumination is applied at 2 Hz, with a pulse duration of 10 ms and peak irradiances ranging from 0.002 to 4.7 mW/mm^2^ on the retina. The light intensity in the projected spot is measured at the cornea and then scaled by the ocular magnification squared, where magnification is defined as the ratio between the sizes of a projected pattern on the retina and in air. VEPs are recorded using the Espion E3 system (Diagnosys LLC, Lowell, MA) at a sampling rate of 2 kHz and averaged over 500 trials. The stimulation threshold is defined as the VEP amplitude exceeding the noise above the 95% confidence interval, which is determined as described in the Supplemental Materials.

## 3. Results

### 3.1 Electrical characteristics of the photodiodes

An example of the dark I-V characteristics for three different pixel sizes is shown in Figure 8a, with the absolute current plotted on a log scale in Figure 8b. The diodes exhibit the expected rectifying behavior. For all three diode sizes, the dark current is in the pA range, the breakdown voltage is around 22 V, and the turn-on voltage at 1 µA is 0.56 V. The single-diode model, described by the equation below, adequately approximates the dark I-V characteristics with ideality factor n=1.5 and a dark saturation current I_0_ = 0.3 pA, as shown in these plots by a black dashed line.

**Figure 8.**
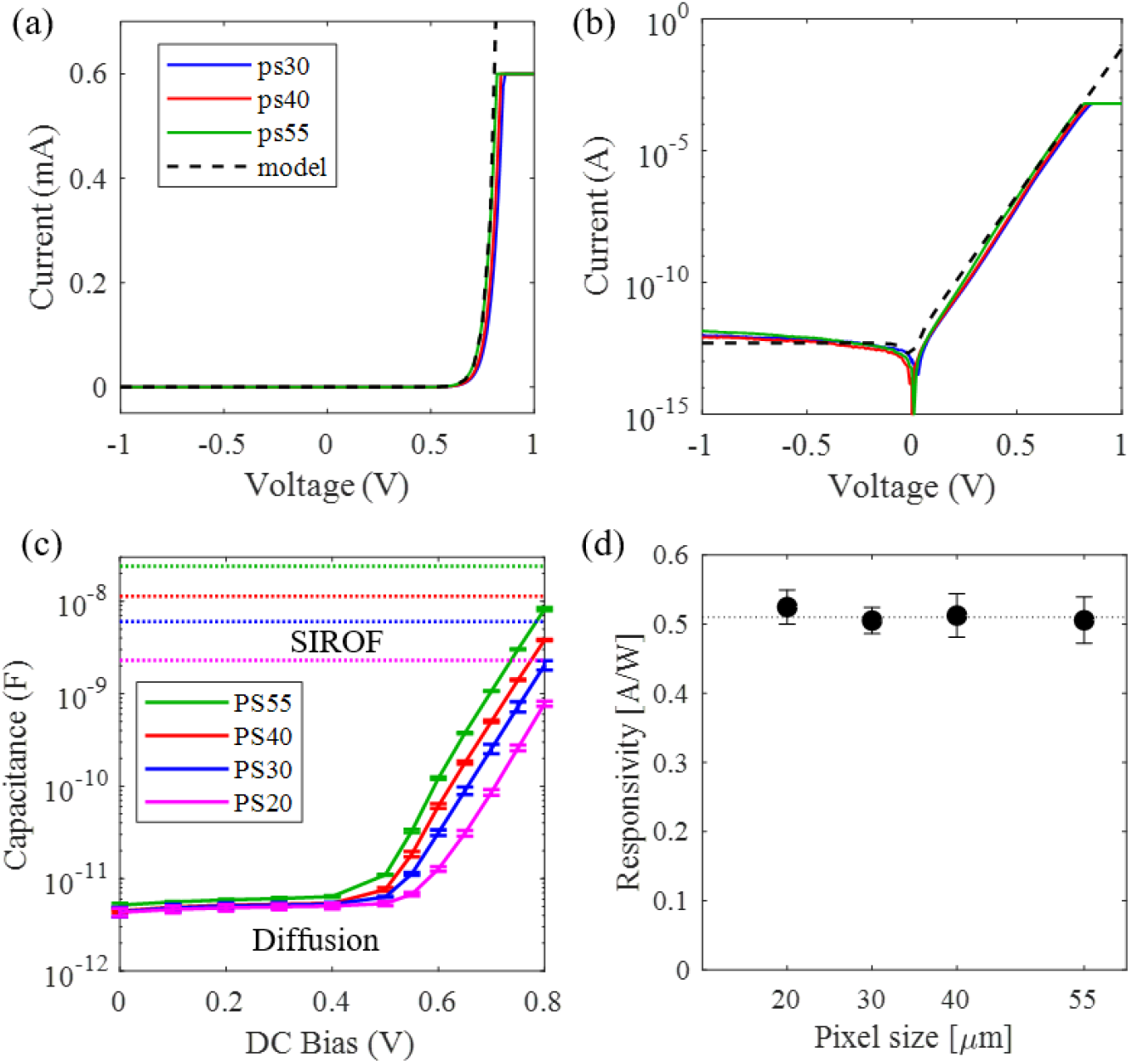
I-V characteristic of diodes with pixel sizes of 55, 40, and 30 µm, shown here on (a) linear and (b) logarithmic scales. (c) Diffusion capacitance as a function of the forward bias voltage. Below 0.6 V, these capacitances are far below the capacitances of the active electrodes, shown by the dash lines. (d) Responsivity of the photodiodes with four different pixel sizes, showing consistently high values independent of size.

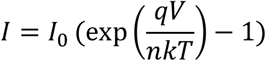

The ideality factor describes the relative contributions of recombination in the depletion region and in the quasi-neutral region. It can be extracted from the fit of the model to experimental I-V curves, and it varies in the range of 1.3-1.6 in the relevant range of voltage (<0.6V). Degradation of the junction quality leads to increase of the ideality factor [43]. Generation-recombination also contributes to an increase in the dark current [44]. Since these diodes are fabricated in a non-commercial setting, we consider the estimated ideality factor and the dark saturation current to be acceptable for our purpose.

Charge accumulation in the quasi-neutral region of the p-n junction under forward bias may affect dynamics of the photovoltaic pixels if it becomes comparable with the charge associated with the capacitance of the electrodes. We measured the diffusion capacitance of the diodes, defined as the voltage derivative with respect to the stored charge. As shown in Figure 8c, the diffusion capacitance increases exponentially with the forward-bias voltage, as expected [45,46]. The forward bias voltage of the Si photodiodes does not exceed 0.7 V under the maximum light intensity in our application (<10 mW/mm^2^). Even at such extremely high bias voltage, the diffusion capacitance is significantly lower than the capacitance of the corresponding SIROF electrodes (Section 3.3.1), by more than an order of magnitude, indicating that the charge stored in the quasi-neutral region is negligible compared with the charge stored in the SIROF electrodes and injected into the medium. Additionally, from the diffusion capacitance at 0.3 V, we estimate the minority carrier lifetime to be 360 μs [45].

### 3.2 Optical response of the photodiodes

The photo-responsivity is a key metric of the photovoltaic pixel performance and directly affects the retinal stimulation thresholds in terms of irradiance. With vertical junctions, we expect responsivity to be independent of the pixel size. As shown in Figure 8d, the measured responsivity is approximately 0.51±0.03 A/W with all four pixel sizes. The inherent limit for responsivity R is described by relating the number of electrons collected to the energy of incident light, thus accounting for losses due to absorption A and internal quantum efficiency η_int_ (R = Aη_int_·*q*/*hf*), where *q* is the charge of an electron, *h* is Planck’s constant, and *f* is the photon frequency. At 880 nm wavelength, the upper limit of Si responsivity is equal to *q/hf* = 0.71A/W. Considering the 85% absorption in our devices, as described in Section 2.5, the upper limit for responsivity for 880 nm radiation is 0.60 A/W. Thus, our measured responsivity of 0.51 A/W corresponds to an internal quantum efficiency of 85%.

### 3.3 Charge injection into the medium

Capacitance of the SIROF layer depends on its thickness [15]. Because of the shadowing effect of the lift-off layer necessary for SIROF patterning, the SIROF coating is thinner at the edge of the electrode than in the center, as can be seen in the FIB cross-section in Figure 9a-c. At the center of this 10 μm electrode, the SIROF thickness is about 300 nm but decreases below 200 nm at the edge. Naturally, such shadowing affects a larger fraction of smaller electrodes. To assess the average capacitance per unit area, we divide the capacitance of the active electrode measured with each of the four pixel sizes by the corresponding electrode area. As shown in Figure 9d, the average capacitance per unit area increases from about 4.5 to about 6.5 mF/cm^2^, with electrode diameters increasing from 8 to 22 μm, corresponding to pixel sizes ranging from 20 to 55 μm. The SIROF thickness on wafer 2 was lower than on wafer 1 by 70 nm. As shown in Figure 10d, the average electrode capacitance of the devices from wafer 2 was, correspondingly, a bit lower.

**Figure 9.**
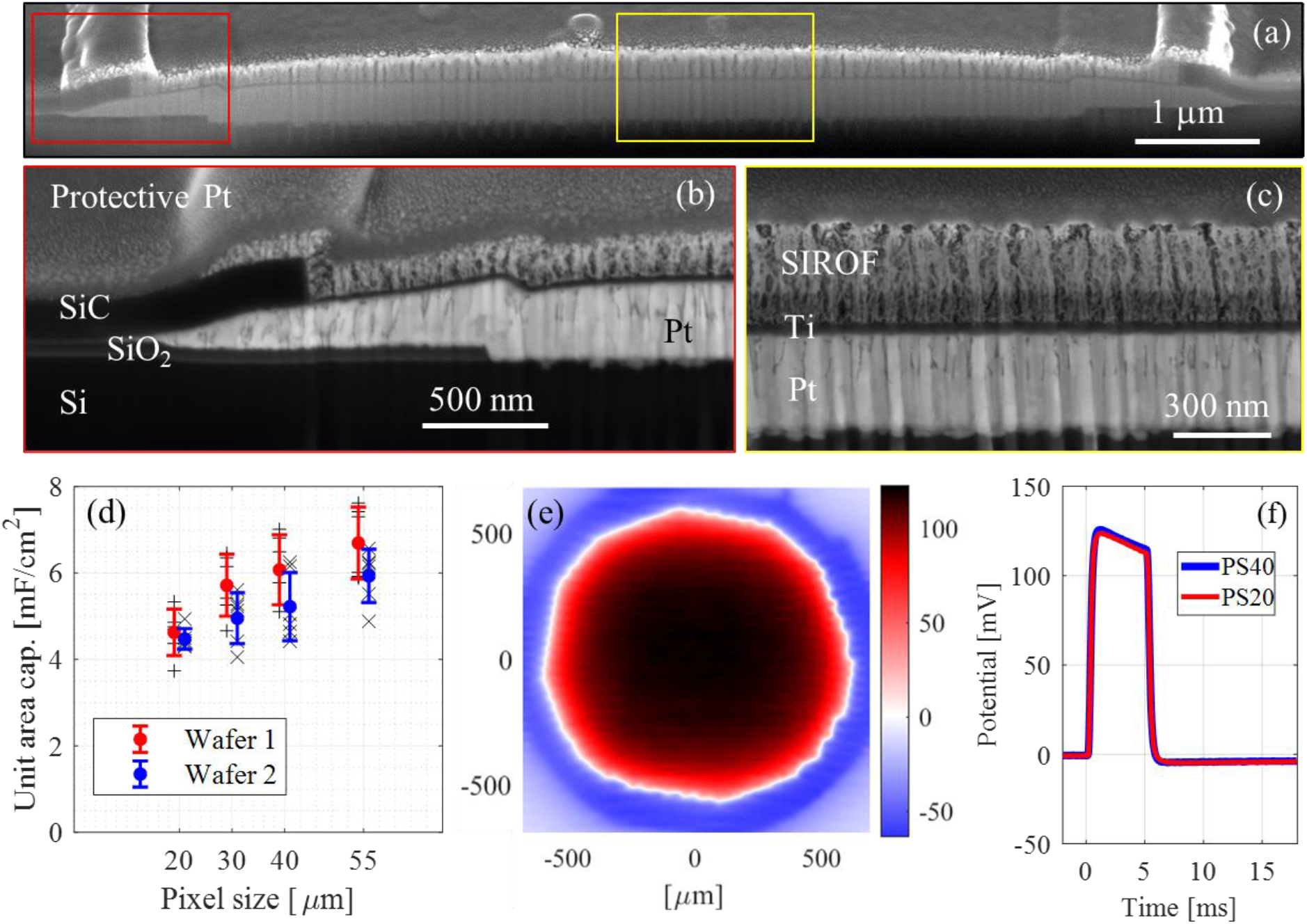
(a-c) FIB cross section of the active electrode. The electrode is composed of Pt, Ti, and SIROF layers, with their thicknesses decreasing toward the edge of the electrode. (d) Average capacitance of the active electrodes decreases with decreasing pixel size due to SIROF thickness non-uniformity. (e) Peak electric potential in the medium 20μm above the device with 40 μm pixels, generated by full-field illumination at 5 mW/mm^2^ irradiance. (f) Waveform of the electric potential measured 20 μm above the centers of the devices is nearly the same with 20 and 40μm pixels. The y-axis label is shared with (e).

**Figure 10.**
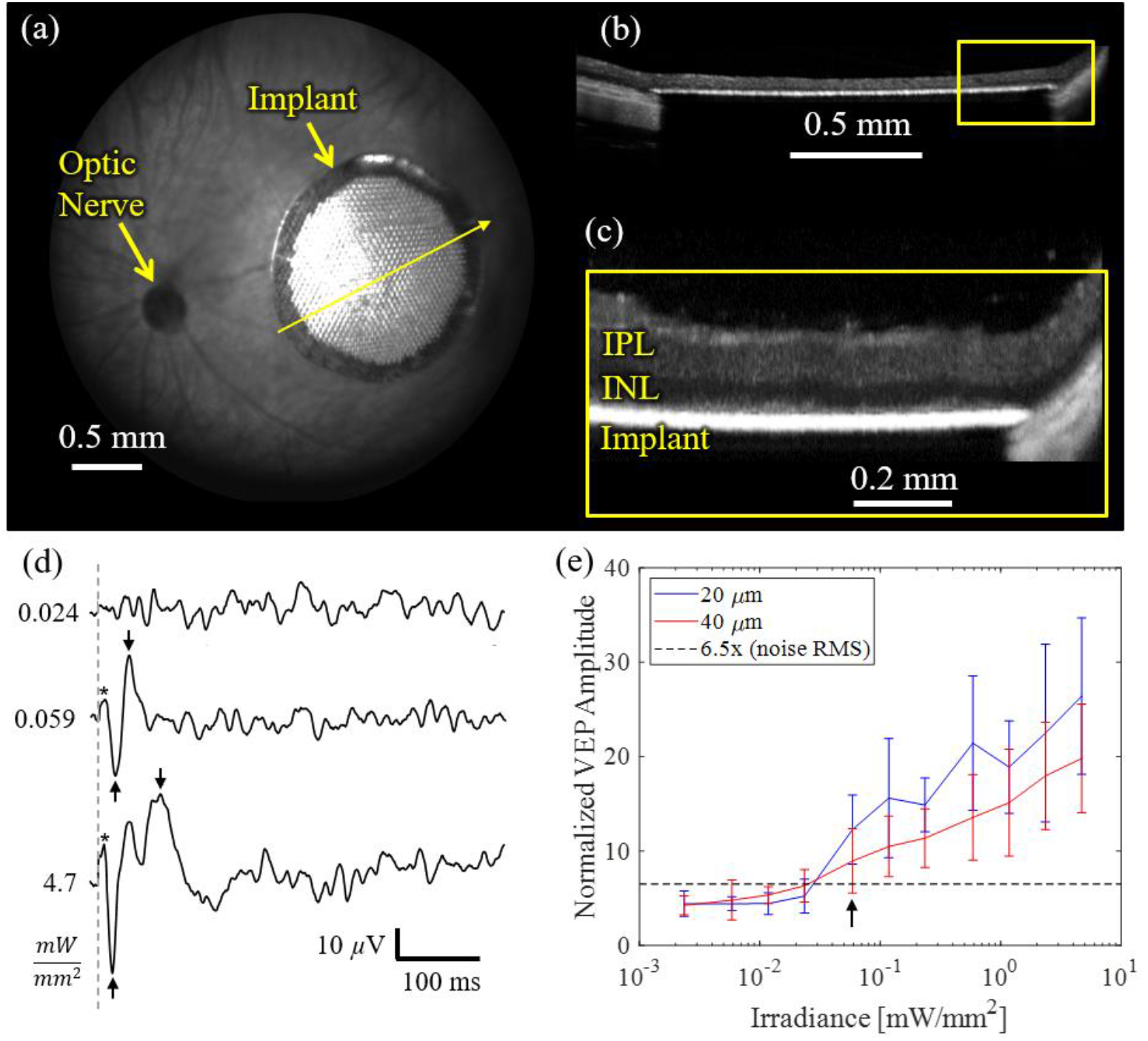
(a) Fundus of the rat eye with a subretinal implant to the right of the optic nerve. (b) OCT image of the implant under the retina and (c) higher magnification demonstrating close apposition of the INL to the implant. (d) Typical VEP in response to the full-field stimulation with 10 ms pulses at various irradiances, repeated at 2 Hz (averaged over 500 trials), with * indicating the stimulation artifact, and arrows pointing at the negative and positive peaks. (e) Average VEP amplitude as a function of the incident irradiance on the retina, with implants having 20 and 40 μm pixels (n=4 per group). The amplitude is normalized to the RMS of noise in each animal. Error bars represent standard deviation. Dash line depicts the 6.5 x (RMS noise) level, which corresponds to the 95% confidence interval.

To assess uniformity of the pixel performance in the array, we mapped the electrical potential induced by the photovoltaic pixels in electrolyte under full-field illumination. Measurements were performed by scanning a micro-pipette in the medium 20 μm above the device with 40 μm pixels. As shown in Figure 9e, all pixels inject current at a similar level, generating a smooth potential distribution, and, as expected, electric potential near the edges of the array is lower due to proximity to the return electrode. Figure 9f shows typical waveforms induced by a 5 ms pulse of NIR light at 5 mW/mm^2^ irradiance, as recorded by a micropipette in the central parts of arrays with 20 and 40 μm pixels. The fact that electric potential in the medium does not depend on pixel size confirms our assumption that a vertically directed electric field eliminates the dependence of penetration depth of the field on pixel size. In the future, this feature will be replicated with honeycomb structures with elevated return electrodes around each pixel.

### 3.4 Stimulation threshold measurements

The animal response to stimulation by the implant was assessed by recording the VEP from the visual cortex, as described previously [47]. Using 880nm light, we projected square patterns of at least 1.5×1.5 mm on the retina to ensure full coverage of the subretinal devices at irradiance ranging from 0.002 up to 4.7 mW/mm^2^. We also verified that there was no response to NIR stimulus when light was projected on the rat retina outside the implant. The VEP amplitude was quantified as a peak-to-peak voltage of the recording within the 10 ms to 200 ms time window. Signals with amplitude greater than noise by 95% confidence (see Supplementary Material) were considered to be responses above the stimulation threshold.

Three example VEP waveforms are shown in Figure 10d, demonstrating a typical cortical noise below the stimulation threshold (0.024 mW/mm^2^), a response just above the threshold (0.059 mW/mm^2^), and the response at the highest irradiance (4.7 mW/mm^2^) in the same animal. The dashed line indicates the beginning of the stimulus, and a 10 ms long small positive peak starting right after the trigger is the stimulus artefact, indicated by the asterisks. VEP amplitude was defined as a peak-to-peak value, which are indicated by the arrows. VEP amplitude varies between animals due to variations in the implant placement, electrode location relative to the visual cortex and the state of anesthesia. To account for the latter two factors in the population averaging, VEP amplitude was normalized by the RMS noise of the cortical signal in each animal. Figure 10e depicts the normalized amplitude of the VEP response, averaged over four animals in each group, having implants of 20 and 40 μm pixels. Error bars represent the population standard deviation. Stimulation threshold with both pixel sizes, indicated by the arrow in Figure 10e, was measured to be 0.057±0.029 mW/mm^2^ (n = 4 for each pixel size), which is about 60 times larger than natural irradiance on the retina (1µW/mm^2^) Above the stimulation threshold, VEP amplitude increases logarithmically with irradiance.

Stimulation threshold of 0.057 mW/mm^2^ with our pixel specifications (ratio of the photosensitive area to the active electrode area = 3.15, photoresponsivity = 0.51 A/W) corresponds to the current density of 0.092 mA/mm^2^ on the active electrode. This result matches the expectation from the modeling described earlier: 0.12 mA/mm^2^. As predicted, it is much lower than with planar bipolar pixels of our previous design, where stimulation thresholds were 0.64, 1.1 and 2.1 mA/mm^2^ with pixels of 70, 55 and 40 μm in width, respectively [12,16].

With 10 ms pulses, current density of 0.1 mA/mm^2^ corresponds to a charge density of 0.1 mC/cm^2^ on the active electrode. With the average capacitance per unit area in the range of 4-8 mF/cm^2^ (Figure 9d) and voltage limit of 0.3V (assuming equal voltage drop on the active and return electrodes), the SIROF electrodes can inject up to 1.2 – 2.4 mC/cm^2^ of charge, which is about 12 – 24 times above the stimulation threshold. Therefore, despite some shadowing at the edges, in principle, electrodes of even the smallest pixels (20 μm) should be able to provide a dynamic range of stimulation exceeding a factor of 10. It is important to keep in mind, however, that the maximum current a photovoltaic pixel can inject is also limited by the voltage drop in the medium, which depends on various factors, including the medium resistivity, sparsity of the projected pattern, and the distance between the active and the return electrodes.

## 4. Discussion

In this study, we developed and validated two important aspects of the photovoltaic implant with small pixels and vertical orientation of electric field in the medium: (1) a new photodiode design with high photoresponsivity independent of pixel size, and (2) retinal stimulation threshold in a vertical electric field as compared to the model and to the previous results with flat bipolar pixels.

(1) Making p-n junctions on the sidewall of pixels allows decreasing the pixel size to 20 μm without a decline in photoresponsivity. To estimate the limits of further scaling the pixel size down, we need to consider several properties of p-n junctions, as well as some practical limits with the current technology. In this discussion, we consider only the photodiode array and neglect the limits on the minimum photosensitive region area per pixel required for stimulation.

a. Trench width: Since the trench walls should be tilted to allow adequate trench filling (about 0.5μm at 30 μm height for the current design), and some width is required at the bottom of the trench to allow for consistent etching of the buried oxide, decreasing the trench width below 1.1 μm would be very challenging.
b. Diffusion width: Diffusion of the phosphorus into the sidewalls of the diodes is determined both by the POCl_3_ doping temperature and time and by subsequent thermal cycles. Careful optimization of the thermal cycles and some compromise in terms of the retrograde boron profile near the top surface of the photosensitive region may allow reducing the final diffused width from 1.5 to 1.0 μm.
c. To control the electric fields in the diode, it is useful to have an undepleted region larger than 1 μm near the center of the pixel. Currently, the depletion width is approximately 1 μm, but it can be reduced to about 0.4 μm by increasing the dopant concentration moderately. Since the pixel pitch should exceed the sum of the width of trench, the two depletion regions, two diffusion thickness and the nondepleted area in the center, the pixel pitch could be decreased to about 6 μm.
d. However, as the pixel size decreases, the photosensitive area becomes increasingly limited by the metal interconnections. Their width is limited by technology and does not scale readily with pixel width. For a lift-off process with sputtered metal, as used for the Pt metallization in this study, the actual metal linewidth is at least 1 μm wider than the patterning layer of the photoresist. Based on the above considerations, the minimum pixel width is probably no less than 10 μm with practically available technology.

Several groups have explored other materials besides Si for retinal stimulation, most notably polymers [48,49], with the claim that arrays made of soft materials can better fit the eye curvature than rigid Si implants. For a sheet of material to deform into a spherical shape, it should be not only bendable, but also stretchable-compressible. This is a very rare property, especially under the forces tolerable by the retina. Our experience has shown that the retina easily settles onto a 30 μm-thick flat subretinal implant of 2 mm in width in human patients, and up to 1.5 mm width in a rat eye, and the implant remains stable after the retinal re-attachment. To cover a larger visual field, multiple tiles can be introduced through the same retinotomy and placed adjacent to each other, as was shown earlier in rabbits [50]. The important advantage of Si, compared to other photovoltaic materials, is its ease of fabrication (readily allowing us to explore vertical junctions), the very high quantum efficiency (about 85% in our pixels) at 880-910 nm and the wavelengths that fit between the photoreceptors sensitivity range on the short end and ocular transparency limit on the long end of this spectral window, as shown in Figure 3a. In our view, this property makes Si a material of choice for photovoltaic restoration of sight in patients who retain some light sensitivity, such as AMD and the majority of RP patients.

(2) Vertical walls of the honeycomb-shaped pixels (above the Si) would direct the electric field vertically, along the axons of bipolar cells, which decreases the stimulation threshold compared to the spherical field emanating from a point source, such as a small single electrode, and even more so compared to planar bipolar pixels with local returns, which generate more localized dipole or even quadrupole fields. In addition, decoupling the field penetration depth (which is set by the wall height) from the pixel width, allows scaling the pixels down to dimensions limited only by the tissue migration. Benefits of the reduced stimulation threshold could be used to either (a) reduce the power of the laser and thus decrease the battery size, or (b) increase the width of the visual field, or (c) increase the dynamic range of retinal stimulation. Future preclinical experiments and clinical studies will show the relative values of these benefits to patients.

Development of the vertical walls for return electrodes is far from trivial: with the wall height of 25 µm (set by the thickness of INL) and its ideal width of 2 µm for 20 µm pixels, the aspect ratio exceeds a factor of 12. Conventional lithography struggles to provide such a high aspect ratio [51,52], and we are evaluating various options. Meanwhile, to demonstrate the effectiveness of directing the electric field vertically, we started the assessment of the smaller pixels and the associated retinal stimulation threshold in a monopolar flat configuration of the array since it generates electric near-field similar to that expected with honeycombs. Under full-field illumination, the field is oriented vertically within the retinal thickness and is independent of pixel size, as shown in Figure 9f. Hence, the stimulation threshold is expected to be similar to that with honeycombs, as shown in Figure 2c. Several differences are expected, however: the deep-penetrating electric field with monopolar arrays may affect the tertiary retinal cells (amacrine and ganglion) differently from the more confined field of bipolar pixels, and this may affect the retinal response to stimulation. In addition, the electric field of the monopolar implant is stronger at the center of the array, while with honeycombs it is much more uniform. Therefore, the stimulation threshold with a monopolar array may be defined by the cells in the center, as opposed to the more uniform response expected with honeycombs. Due to a strong cross-talk expected with a monopolar array, our measurements here are limited to full-field stimulation, and not attempting to measure visual acuity using alternating gratings.

## 5. Conclusions

Decrease in size of the photovoltaic pixels for a retinal prosthesis necessitates a transition from planar to 3-dimensional geometry of the electrodes and of the p-n junctions. We demonstrate that diodes with vertical junctions provide very high quantum efficiency (85%), independent of pixel size, at least down to 20 μm. We also show that vertical electric fields practically do not change with the pixel size, and therefore implants with all pixel sizes provide the same retinal stimulation threshold, which is much lower than with bipolar pixels. The next step will be fabrication of the honeycomb return electrodes on top of such photodiode arrays and testing their performance in-vivo.

## 6. Acknowledgments

We would like to thank Jerome Pons from JH Technologies (www.jhtechnologies.com) for their help with sample preparation and SEM imaging of the cross-sectional view of the pixels shown in Figure 5b. Studies supported by the National Institutes of Health (Grants R01-EY-027786, P30-EY-026877), the Department of Defense (Grant W81XWH-19-1-0738), AFOSR (grant FA9550-19-1-0402), Wu Tsai Institute of Neurosciences at Stanford, and unrestricted grant from Research to Prevent Blindness. Photovoltaic arrays were fabricated at the Stanford Nano Shared Facilities (SNSF) and Stanford Nanofabrication Facility (SNF), which are supported by the National Science Foundation under award ECCS-1542152. KM was supported by a Royal Academy of Engineering Chair in Emerging Technology. Part of this work was also performed at the Marvell Nanofabrication Laboratory at the University of California-Berkeley and at the Lurie Nanofabrication Facility at the University of Michigan.

D.P. and T.K. are consulting for Pixium Vision. D.P.’s patents related to retinal prostheses are owned by Stanford University and licensed to Pixium Vision. All other authors declare no financial interests.

