## Supplemental Materials for "Vertical-junction Photodiodes for Smaller Pixels in Retinal Prostheses"

### Supplementary Materials

Retinal migration into the honeycombs

Cells of the inner nuclear layer (INL) in the degenerate retina migrate into the subretinally-implanted honeycomb array [16,17,18], while bipolar cells retain their axons in the inner plexiform layer. Images in Figure S1 were obtained from confocal fluorescence microscopy of the whole-mount retina with the 40 μm-pixels array 6 weeks post-implantation in a 6 months-old RCS rat.

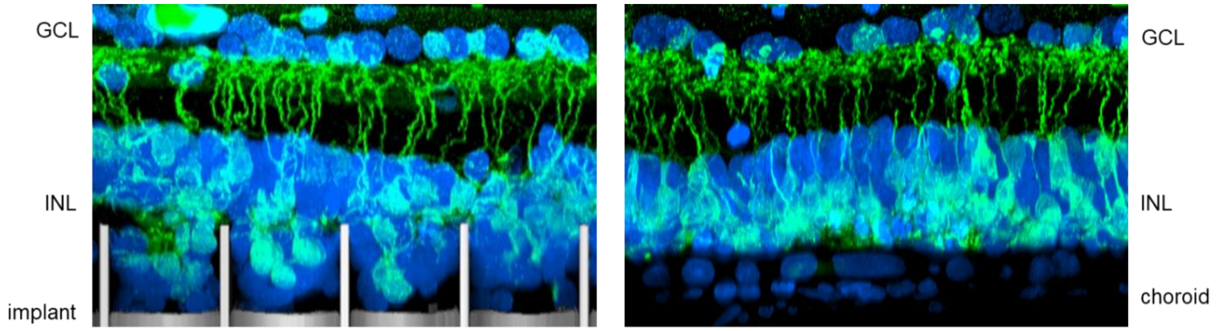


**Figure S1**: *Retinal cells migrating into the honeycomb implant 6 weeks post-op (left) and the retina away from the implant in the same eye (right) of the RCS rat. Blue stain is DAPI, marking all nuclei, and green – PKCα, marking the cone bipolar cells.*

Junction dopant profiling

In order to obtain the dopant profiles shown in Figure 5c, we used TCAD Sentaurus simulation and nanoscale secondary ion mass spectroscopy (nanoSIMS). To simulate gaseous phase phosphorus diffusion in the Sentaurus modeling, phosphorus-doped oxide was deposited onto the side walls at the solubility limit of phosphorus at 1000 $^{o}$C, diffused for 30 min at 1000 $^{o}$C and then removed. We then simulated diffusion of the phosphorus from silicon into the polysilicon filling of the trenches during the subsequent heat treatments. For the nanoSIMS imaging of the phosphorus doping concentration within the sample, we polished flat a sample after all its doping and heat treatments. To obtain the line profile shown in Figure 5c, each column was integrated to provide an average value as a function of distance from the trench.

Photosensitive area calculation

Due to inherent bias during the fabrication procedure, the achieved metal pattern widths do not perfectly match the mask patterns. In particular, the Ti/Pt metal stack is deposited by sputtering onto bilayer resist, which results in sputtering of the metal into the undercut of the liftoff resist layer. This results in some thin 'tail' to the metal at the pattern edges, as seen in Figure 9b, partly due to the shadowing of the resist. To calculate the amount of light reaching the Si for estimation of the photoresponsivity, we approximated the thickness of the tail end of platinum as a linear slope, and knowing the Pt refractive index at 880nm, calculated transmission of light through the thin platinum in that area.

Measurement of the SIROF capacitance

For assessment of the SIROF electrodes and the photovoltaic pixels, we fabricated various test structures on the same wafers with the photovoltaic arrays. Figure S1 shows a test structure which contains 8 photovoltaic pixels, two of each size, on the right side, with leads connecting the active and the return electrodes to metal pads on the left side. The leads, around 2.5mm in length, are passivated by SiC.

A drop of phosphate buffered saline (PBS) solution was placed on the pixel end, while the metal pads in contact with the measurement probes were not wetted.

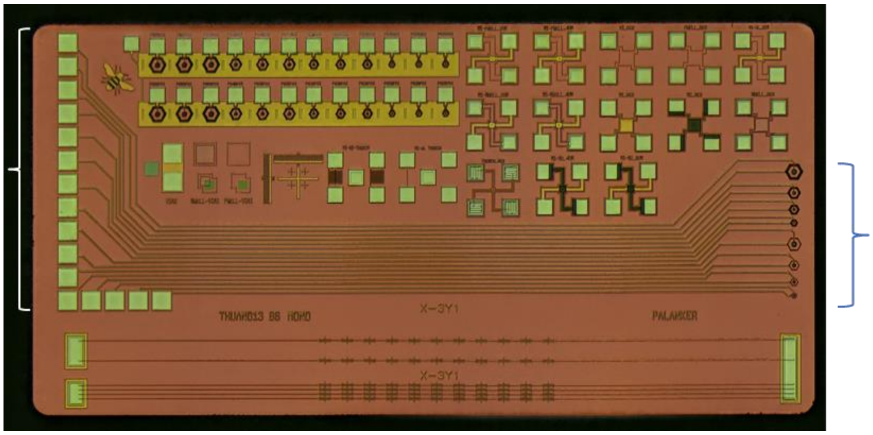

**Figure S2**: *Micrograph of a test structure having the active and return electrodes in each pixel on the right connected to the pads on the left, shown by the brackets.*

To measure the impedance spectrum between a pair of SIROF electrodes, we used a 2-electrode setup at 0V bias. The complication of introducing a bulky reference electrode in such a limited volume of electrolyte could be avoided because the impedance spectrum of a SIROF electrode is independent of the voltage bias within the water window [40]. The impedance spectroscopy was performed with a potentiostat (Interface1010E, Gamry Instruments, Warminster, PA), and the results were fit to the circuit model of a simple Randles cell (Figure S2) [41], from which the capacitance per unit area was calculated based on the measured area of SIROF coverage on each electrode. The fit was implemented using the method of least squares on the relative error, as described in Section 3.3.2.3 of [ref 1]. N=6 electrodes were measured for each pixel size.

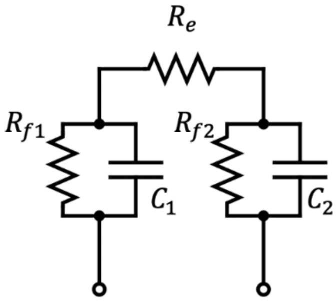


**Figure S3**: *Circuit diagram of a simple Randles cell model the impedance spectra were fit to.*

<u>Determination of the stimulation threshold</u>

Visual cortex exhibits electrical activity even in the in the absence of the visual stimulation, and its level in rats, as recorded in our system, corresponds to approximately 10 μV after 500 averages. To define a VEP response to a visual stimulus with confidence, it must exceed the background activity above the confidence interval. The following procedure was applied to calculate the 95% confidence interval (CI) for a VEP amplitude exceeding the noise: We took a 250-seconds long recording without a stimulus, randomly picked 500 start times in the recording and took the 200 ms following each start time as a segment. Averaging over these 500 segments, we took the peak-to-peak amplitude of the resultant averaged segment as noise. By repeating the random segmentation for 10,000 times on the same 250s recording, we computed the 50th and 97.5th percentile of the resultant 10,000 peak-to-peak amplitudes, and determined the median and 95% CI, as shown in Figure S3.

VEP amplitude varies between the animals due to variations not only in the implant placement, but also in electrode location relative to the visual cortex and in the state of anesthesia. Therefore, for the population averaging, VEP amplitude was normalized by the RMS noise of the cortical signal in each animal. To visualize the stimulation thresholds for multiple sets of measurements, we found the VEP amplitude corresponding to the 95% CI for each set, normalized this amplitude to the RMS of noise, and averaged across the animals. The resulting average normalized signal-to-noise level corresponding to 95% CI is 6.5x RMS of noise, as shown in Figure 10e.

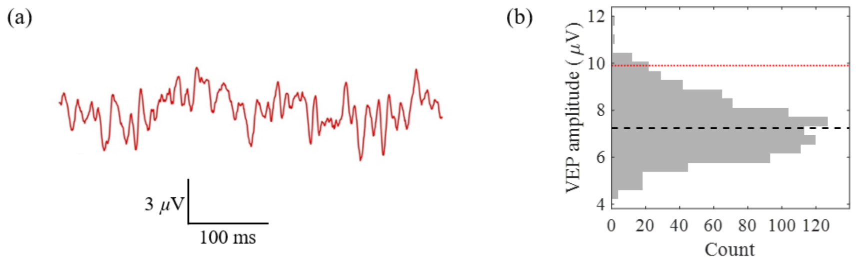


**Figure S4**: *(a) An example VEP recording without any stimulus, averaged over 500 trials. (b) An example of the noise distribution in VEP recordings and determination of the 95% confidence interval (red dotted line) and the 50% confidence interval (black dash line).*
